# Inspecting abundantly expressed genes in male strobili in sugi (*Cryptomeria japonica* D. Don) via a highly accurate cDNA assembly

**DOI:** 10.1101/2020.04.21.054320

**Authors:** Fu-Jin Wei, Saneyoshi Ueno, Tokuko Ujino-Ihara, Maki Saito, Yoshihiko Tsumura, Yuumi Higuchi, Satoko Hirayama, Junji Iwai, Tetsuji Hakamata, Yoshinari Moriguchi

**Author notes:** Corresponding author (SU).

## Abstract

Sugi (*Cryptomeria japonica* D. Don) is an important conifer used for afforestation in Japan. The field of functional genomics is rapidly developing. The genomics of this gymnosperm species is currently being studied. Although its genomic size is 11 Gbps, it is still too large to assemble well within a short period of time. Transcriptomics is the one another approach to address this. Moreover, it is a necessary step in obtaining the complete genomic data. Here we designed a three stages assembling workflow using the *de novo* transcriptome assembly tools, Oases and Trinity. The three stages in transcriptomics are independent assembly, automatic and semi-automatic integration, and refinement by filtering out potential contamination. We found a set of 49,795 cDNA and an equal number of translated proteins (CJ3006NRE). According to the benchmark of BUSCO, 87.01 % were complete genes, including very high “Complete and single-copy” genes–78.47%. Compared to other full-length cDNA resources, the extent of the coverage in CJ3006NRE suggests that it may be used as the standard for further studies. When two tissue-specific libraries were compared, principal component analysis (PCA) showed that there were significant differences between male strobili and leaf and bark sets. The highest three upregulated transcription factors stood out as orthologs to angiosperms. The identified signature-like domain of the transcription factors demonstrated the accuracy of the assembly. Based on the evaluation of different resources, we demonstrate that our transcriptome assembly output is valuable and useful for further studies in functional genomics and evolutionary biology.

## Introduction

*Cryptomeria japonica* D. Don (Japanese cedar), also known as sugi in Japan, is a large evergreen conifer tree species. Because of its fast growth and its adaptation to most environments in Japan, it has been an important material for the forestry industry. After World War II, sugi plantations have increased to 42 % of Japan’s artificial forests [1]. Thus, the need for breeding better tree varieties is one of the main reasons to attain more knowledge of sugi genomics. Other motivations include medical and other economic reasons. Its pollen led to severe allergy in about 25% of the Japanese population [2]. Replacement by sugi with sterile pollen is a possible solution to the problem, but it is not easy to implement and may take a long time [3]. Fortunately, 23 genetically sterile male trees have already been identified [4]. Four loci (*MS1, MS2, MS3*, and *MS4*) responsible for male sterility were located in four different linkage groups (LG9, LG5, LG1, and LG4, respectively) [5]. In addition, male sterility is caused by recessive alleles. Recent advances in technology are available to reveal more details on the genetics related to these loci. To more precisely identify the genetic variation or the genes related to male sterility, a functional genomic study via transcriptomics is a logical approach [6, 7], though a comprehensive genomic data of sugi is not currently available.

In gymnosperms, study on functional genomics is difficult because of the long life span of trees and the large genome size. For instance, the genome sizes of Norway spruce (*Picea abies*) [8], white spruce (*Picea glauca*) [9, 10], loblolly pine (*Pinus taeda*) [11], ginkgo (*Gingko bioloba*) [12], and sugi (*Cryptomeria japonica*) [13] are 20 Gb, 20.8 Gb, 20.15 Gb, 11.75 Gb, and 11 Gb, respectively. So, far, only four gymnosperms have had their assembled genome sequences published. That is, the Norway spruce [14], white spruce [9, 10], loblolly pine [11], and ginkgo [12]. The genomic sequences have been assembled from long length of contigs or scaffolds. Decoding the genome using appropriate annotations is essential for determining the functions. There has been an increase in the use of the emerging annotation software— “MAKER” and “MAKER-P” for plant species [15]. Genomic information from assembled genome sequences, RNA, and protein data can be combined and annotated. In other words, regardless of the quality of the genomic sequence, abundant RNA or protein data is required for the annotation process. Thus, even where a complete genome sequence is unavailable, transcriptome analysis is needed for a functional genomic study [8, 16-20].

Many assembled expressed sequence tags (EST) for sugi have been published in the public databases [21, 22] before the availability of high throughput sequencing technology—such as next generation sequencing (NGS). This was sufficient for functional genomic studies for certain specific purposes. However, NGS has proven to be greatly beneficial to the advancement of functional genomic research due to its increased yields, reduced unit price, and multiple analysis tools available [7, 23]. Pacific Biosciences (PacBio) is a single-molecule real-time long-read isoform third-generation sequencing tool. It provides a reasonable alternative to harvesting the full-length cDNA. The length of a single read and the high speed were unprecedented [24]. However, errors still needs to be accounted for by hybrid genome sequencing assembly [25]. Choosing a suitable assembly tool is another issue in processing in the *de novo* assembly of transcriptome data [26]. Transcriptomic integration is another option, which was performed well in *Abies sachalinensis*, another conifer species [27].

Fundamental information on the transcriptome is an essential step for future work; we aim to construct a high-quality cDNA assembly on sugi. Here, we independently assembled the transcriptome data of 10 different genome accessions of sugi. These were then integrated with half-manual integration using the EvidentialGene software [28]. To find and identify the male strobili specific sterility genes, we had the 10 RNA-Seq libraries of uneven runs. In reference to the benchmark testing and coverage of to- and by-different cDNA sources, we have high confidence in our assembled cDNA.

## Results

The integrated sugi EST library containing 49,758 transcripts, has been constructed by a series of manipulation on the 10 RNA-Seq libraries. We joined two RNA-Seq *de novo* assembling tools (Oases [29] and Trinity [30]); and used two methods to assemble and to integrate the transcriptomic sequence with the RNA-Seq libraries (automatic assembly using EvidentialGene [28] and half-manual assembly in multiple steps). We evaluated our library with the benchmark tool using the built-up core reference database; and we compared our integrated EST library to the other cDNA resources. The expressional result (Figure S2) showed the highest effect by the tissue (Table 1) compared to the pedigree of samples (Figure S1). The purpose of these evaluations is to assess the quality of the transcriptome assembly output from this method.

**Table 1.**
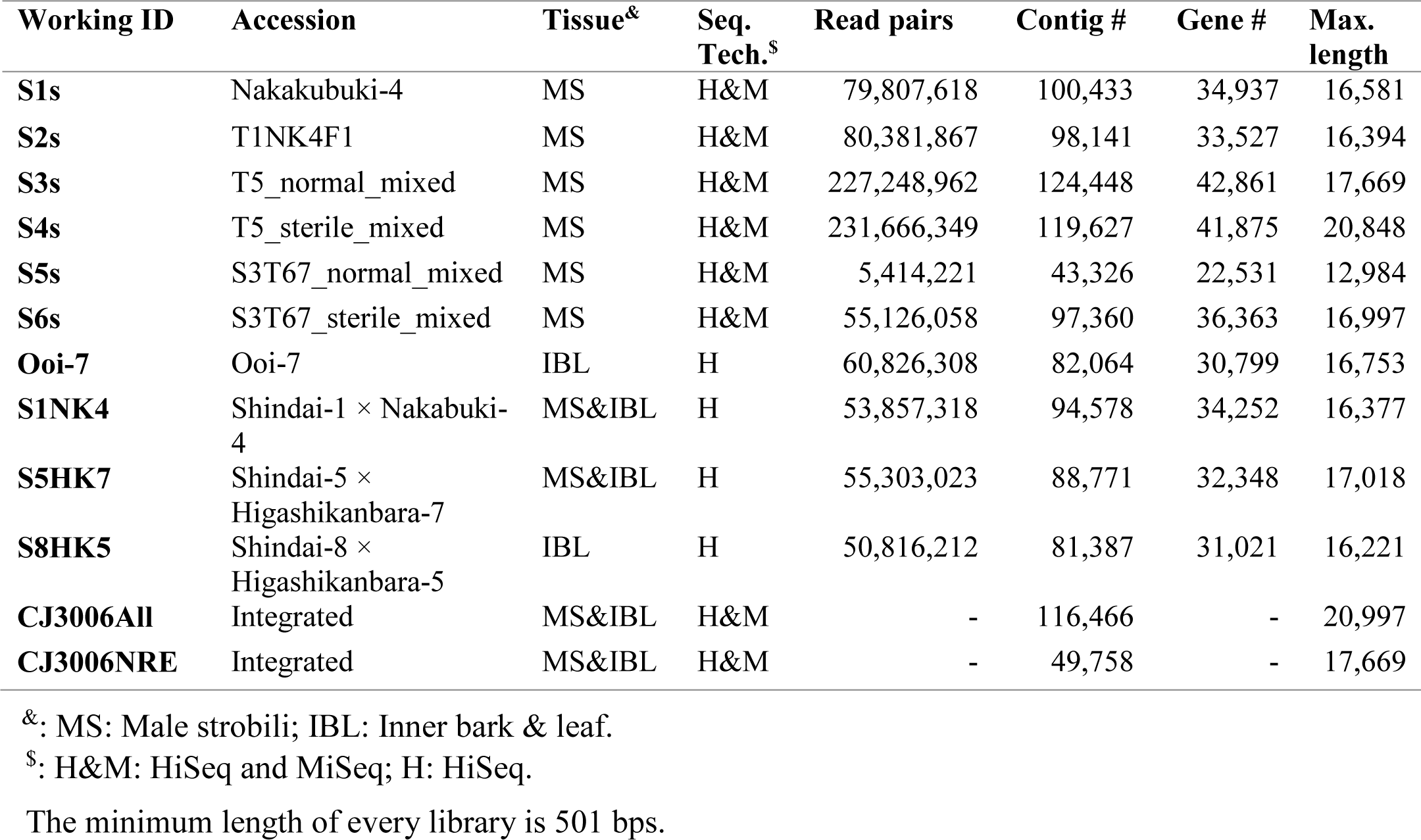
The statistics of raw reads and mapped read

### Assembly and annotation of EST library

There were 116,466 and 49,758 contig sequences in CJ3006All and CJ3006NRE, respectively. The maximum cDNA length of CJ3006All and CJ3006NRE were 20,997 and 17,669, respectively. The N50 statistic of CJ3006All and CJ3006NRE were 1,256 and 1,819, respectively. The basic number of each assembled library is presented in Table S1. In Trinity, the number of contigs per library ranged from 43,326 (S5s) to 124,448 (S3s). In Oases, the number of contigs per library ranged from 34,303 (S5s) to 105,184 (S3s). Obviously, the number of contigs was affected by the number of reads per library (Table 1).

There were a total of 31,678 and 47,968 genes out of 49,758 to which we assigned a functional annotation using InterProScan [31] and EvidentialGene [28], respectively. It is difficult to identify the real gene isoform from the assembled transcriptomes without genomic sequences. Although 17,079 gene isoforms (Table S2) of the 49,758 can be used as representatives, we used all 49,758 genes to perform the following analysis.

In total, 1,291 genes related to transcription factors have been identified by Pfam [32] annotation as in the list of transcription factors (Table S2). Within them, 974 genes were considered as unigenes without isoforms (Table S2).

Using RepeatMasker [33], the number of transposable elements within CJ3006NRE was estimated to be 7,029 and 2,282 for retrotransposon (Class 1) and DNA transposon (Class 2), respectively. Repetitive sequences made up about 4.1 % of the whole cDNA sequences in CJ3006NRE (Table S3). The majority of this was LTR elements, forming about 2.54% of nucleotide bases in the total length of CJ3006NRE. However, only 3,197 and 772 were with confidence after filtering out ones with low coverage rate (<20%) and too short length (< 200 bps) (data not shown). Although retrotransposons may account for the large size of the sugi genome, most of them may be silenced in the collected 10 RNA-Seq libraries.

The cDNA and translated protein sequences were uploaded onto the ForestGEN database (http://forestgen.ffpri.affrc.go.jp/CJ3006NRE/clusterList with username and password, which will be freely accessible after acceptance of this manuscript). The metadata and annotation are presented in a Supplementary Excel file (Table S2).

### The benchmark of assembly

The coverage of the assembling result of the sugi EST library (Fig 1) was estimated by BUSCO (version 3.0.2) [34] and its reference database, embrophyta_odb9. In comparison to other assembled results from single libraries, S1s to S8HK5, the ratio of missing parts (to the reference database) was lower in CJ3006NRE, 10.76%. Furthermore, completeness, especially the “Complete (C) and single-copy (S),” was clearly higher (78.47%) than all other libraries (41.88% to 49.24%). In general, the benchmark of CJ3006NRE is higher than any single assembled library.

**Fig 1.**
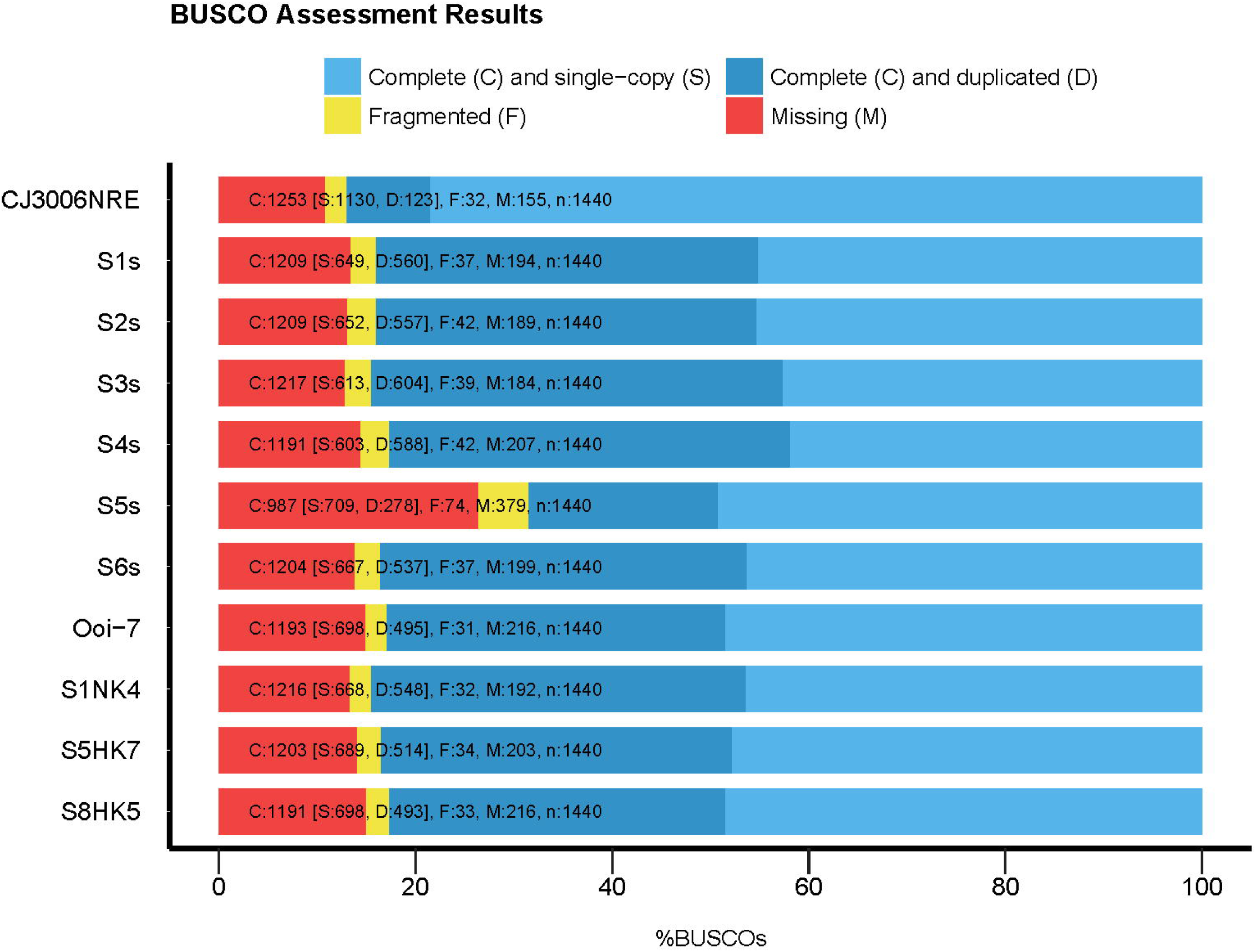
The benchmark of the assembled contigs. The y-axis shows the samples listed in Table 1. The reference database for BUSCO (v 3.0.2) was “embrophyta_odb9.” with 1,440 core genes in total.

### The coverage of different full length cDNA

For estimating the advantage of the RNA-Seq technology and integration method used in this study, we compared the coverage between CJ3006NRE and two other cDNA sources, full-length cDNA by Sanger sequencing and full-length cDNA by ISO-Seq results in our laboratory.

The full-length cDNA library was retrieved by using the keywords “Cryptomeria japonica full-length” in the NCBI-Nucleotide database on 13th September, 2018. In total, 23,111 nucleotide sequences were retrieved, downloaded, and formatted as BLAST databases, called “CJ_FLcDNA.”

Using the “pbtranscript-tofu” analysis suite, there were 56,399 transcripts clustered from three ISO-Seq runs. Within them, 9,352 transcripts were classified as a high-quality subset, called “ISOSeq0215hq”.

For screening on the BLAST result, we merged the High-scoring Segment Pairs (HSP) of the same query-subject pair. Then, we separately calculated the coverage to query and to subject. Thus, we calculated the coverage to the “query” sequence and to the “subject” sequence for each query-subject pair. In Table 2, we only counted the sequences where the coverage was over 75% of the length. Every row of Table 2 shows that using CJ3006NRE can cover a higher ratio of another cDNA library than vice versa. Thus, using CJ3006NRE to present both cDNA resources would have better overall coverage.

**Table 2.**
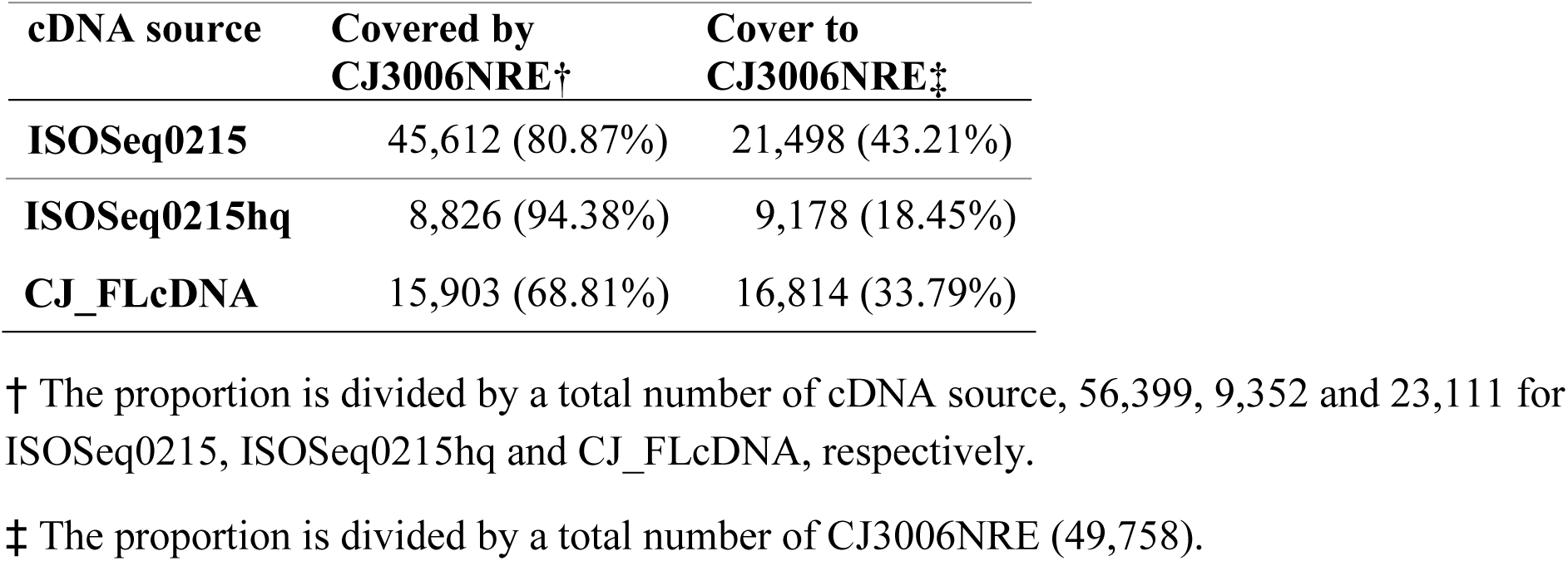
The mutual coverage (more than 75%) between CJ3006NRE and other cDNA sources

### The mapping rate of RNA-Seq reads

The usage of the reads was counted based on the statistics of the mapping file. The mapping rate (%) was calculated by dividing the number of unmapped reads by the total number of reads of each library, and then subtracting from one hundred.

Since the integration process has been through several steps, we used mapping (usage) rate to reveal how much information by sequencing was lost. Using CJ3006NRE as the reference, the usage rate ranged from 91.99 % to 96.49% (Fig 2; Table S4). Theoretically, the usage rate against the contigs assembled from the querying reads should be one hundred percent. However, we could only have 96.27% (S5s) to 99.04% (S4s). After the integration, however, almost all mapping rates reduced by around 0.87% (S3s) and 2.09% (S1s), except for S5s, which increased by 0.74%. By filtering out the non-eukaryote assemblies (i.e., contamination), CJ3006NRE was attained; the mapping rate reduced by another 1.58% (S4s) to 3.35% (S1NK4). Thus, there was only a low number of reads that have been discarded by the integration and filtering processes.

**Fig 2.**
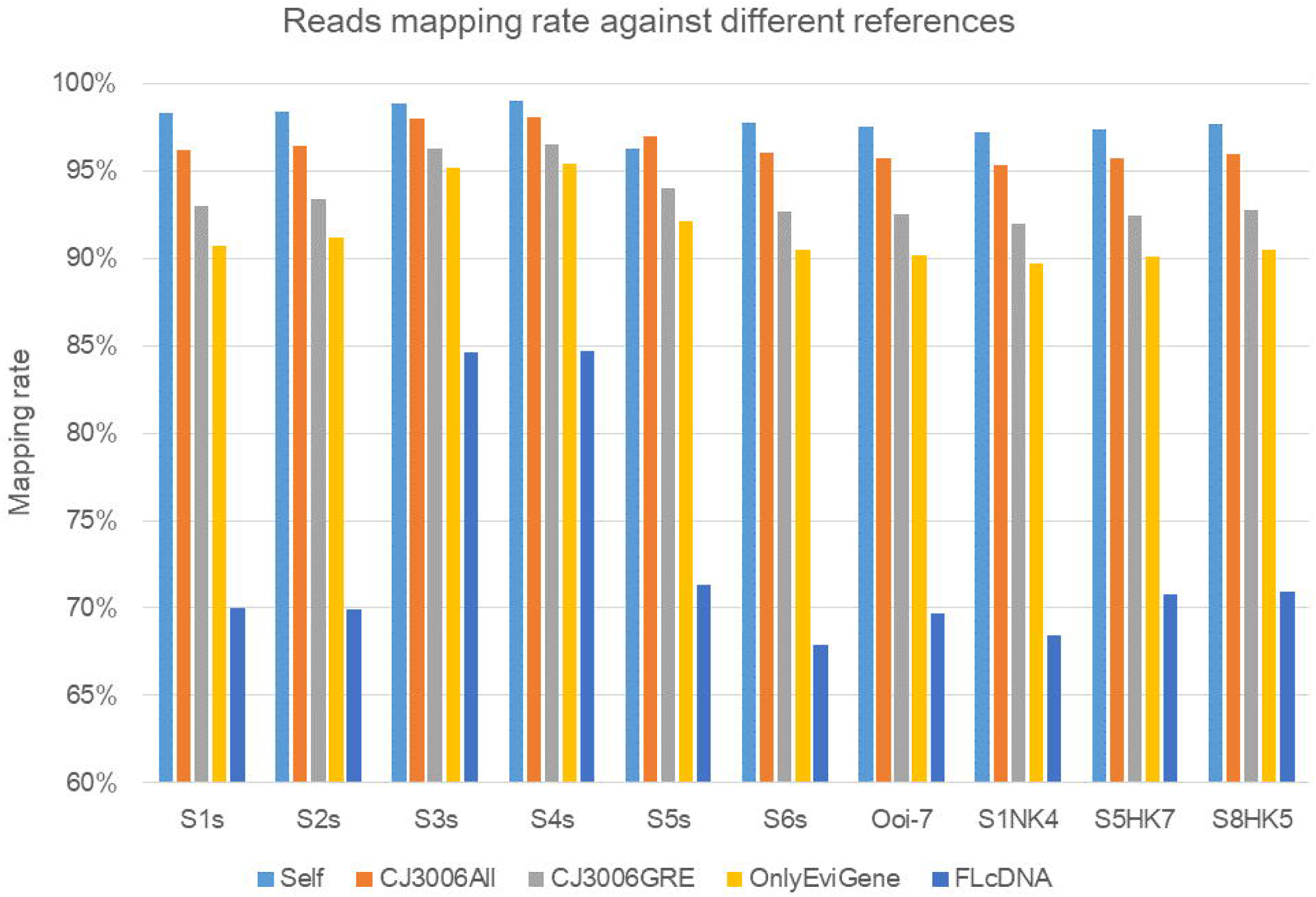
The mapping rates from different references. The reads mapping rate was estimated by samtools on the mapping file (BAM). The X-axis means the source of raw reads. The five colors represent the different reference contigs. Self: the reference was the contigs assembled by the same library of raw reads; CJ3006All: the contigs set of CJ3006All without filtering; CJ3006NRE: the contigs set of CJ3006NRE, the CJ3006All without matching to any non-Eukaryote in NCBI-NR; Only EviGene: CJ3006NRE without manual integrated set; FLcDNA: the full-length cDNA set download from NCBI on 13^th^ September, 2018. The Y-axis is the ratio of mapped raw reads against the reference contigs.

For determining how much reads was enriched by manual integration, the column “Only EviGene” was calculated (Table S4). It showed enrichments of 1.09% (S4s) to 2.31% (S1NK4). Compared to the added contigs number of the manually integrated part, 7,580 contigs, which is about 15.23% of CJ3006NRE, only a small amount of enrichment was given by the manual integration.

By mapping to full-length cDNA from NCBI, we determined how much the RNA-Seq data covers the full-length cDNA, particularly for the cDNA collected from the same plant organ - the male-strobilus. The result showed a distribution of 67.92% (S6s) to 84.69% (S4s) (Table S4). The two highest mapping rate was contributed by two libraries with the highest numbers of reads, S3s and S4s. This suggests the full-length cDNA from NCBI only covers less than 85% of the total RNA, if we assume RNA-Seq as the total RNA.

### Differential expression

The 10 RNA-Seq libraries could be classified into three groups, according to the type of tissue sample collected: 1.) All from male strobili; 2.) All from the inner bark and leaf; and 3.) Mixed with male strobili, inner bark, and leaf materials. Before looking at the differential expression between any certain pair of groups or libraries, principal component analysis (PCA) and sample heatmap were used to reflect the characters of the libraries. The two libraries collected from only inner bark and leaf tissues (Ooi-7 and S8HK5) were very different from other libraries (Fig 3). In PCA, either the first or second principal component divided the libraries from bark and leaf far from the accessions from male strobili (Fig 3a). Although most of the samples were collected from male strobili, bias during sampling or assembling may occur. In Fig 3b, the darker blue indicated a higher dissimilarity between two accessions, as indicated by the X-axis and Y-axis. With the exception of “Ooi-7 against S8HK5”, almost all other accessions against Ooi-7 or S8HK5 showed higher dissimilarity (> 0.1, darker color). This supported the previous PCA result in one-to-one comparisons.

**Fig 3.**
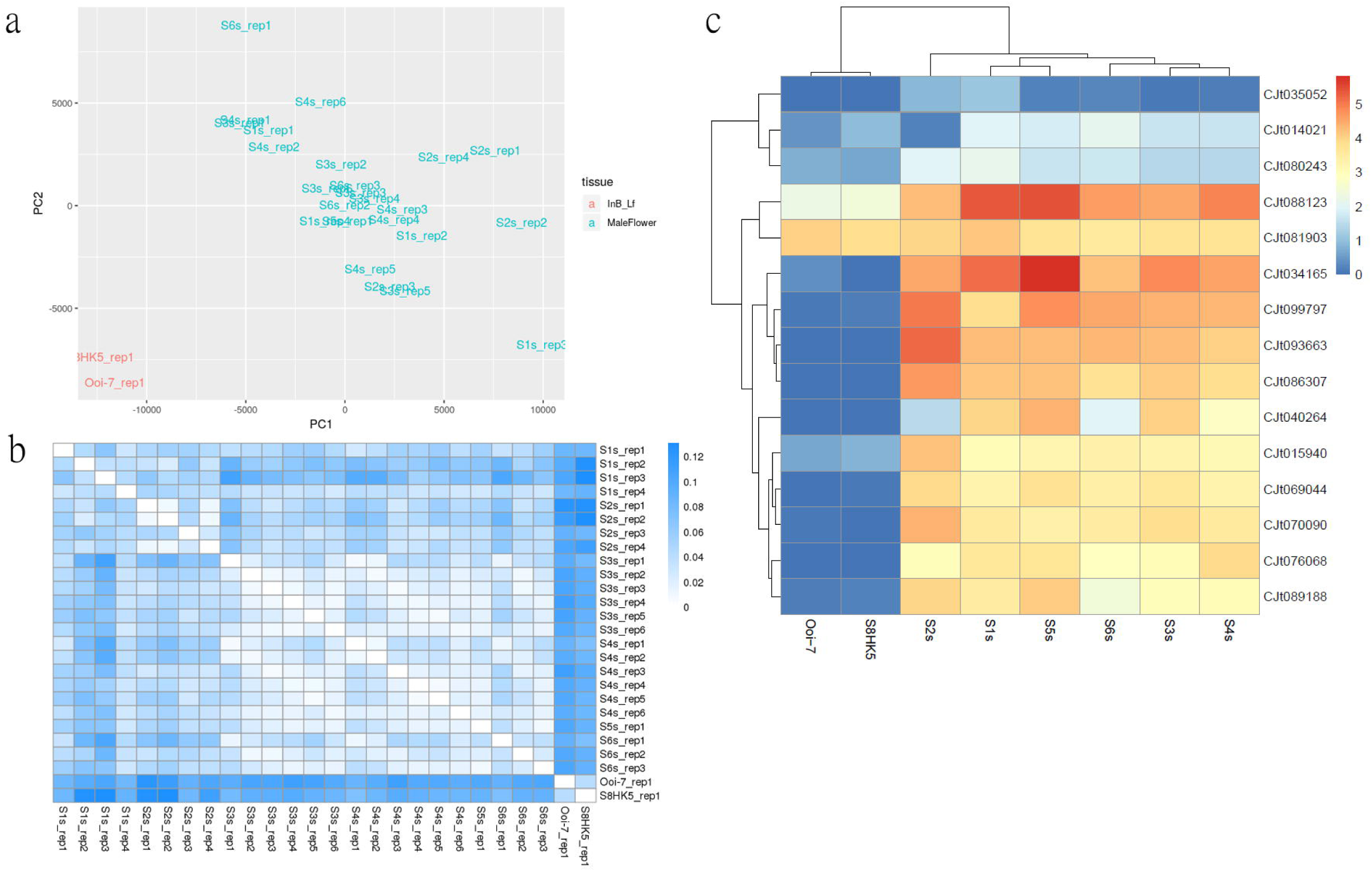
Assessment of the gene expression data. (a) The PCA result of each run. The green color indicates samples collected from male strobili. The red color indicates samples from bark or leaf tissue. (b) Paired divergence heatmap among each run. The darker blue indicates higher divergence. (c) The heatmap of gene expression of potential genes downstream from MYB80s

Two runs of the S1 assembly (i.e., S1s_rep2 and S1s_rep3) showed high dissimilarity to several other runs collected from different times. The explanation for the dissimilarity of these two runs to the others is uncertain. However, two other runs of S1 (i.e., S1s_rep1 and S1s_rep4) supported the similarity of S1s to those from the male-strobilus libraries. Nonetheless, the PCA result showed S1s_rep1, S1s_rep2, and S1s_rep4 were all clustered with the other male-strobilus accessions. Thus, the divergence did not have large effects on the overall clustering result.

The expression between the male-strobilus set (library S1s to S6s) and inner-bark-and-leaf set (library Ooi-7 and S8HK5) were significantly different in 7,776 genes (P-value < 0.05) (Table S5). Within these, 4,471 are upregulated in the male strobili with a fold change ranging from 2^0.33^to 2^10.28^. The fold change range for the rest of 3,305 down-regulated genes was from 2^−6.41^to 2^−0.33^. The heatmap showed different patterns among different tissue sets (Figure S2).

There was a total of 377 transcription factors within the 7,776 genes with significantly different expression (Table S5). Of these, the three largest gene families were MYB DNA-binding, HLH, and AP; which consisted of 56, 43, and 42 genes, respectively (Table S6). Interestingly, the trend among these three gene families and between the two tissue types is the same. The number of transcription factors in the male strobili was about 3 times higher than leaf or inner bark, as well as higher levels of gene expression. Clearly, the activities occurring in male strobili require higher gene regulation.

The downstream genes corresponded to AtMYB80 (At2g47040, named VGD1), are a glyoxal oxidase (At1g67290, named GLOX1), and an A1 aspartic protease (At4g12920, named UNDEAD) in *Arabidopsis*. The potential orthologs in CJ3006NRE are VGD1: CJt093663 and CJt014021; GLOX1: CJt015940 and CJt035052; UNDEAD: CJt080243 and CJt088123. Since the exact orthologs are unclear, the top two subjects were selected according to the HSP, which covered over 80% of the length of query sequences. Within each potential orthologs group, only one gene showed significantly higher expression in MF than in IBL. They are CJt093663, CJt035052, and CJt080243 for VGD1, GLOX1, and UNDEAD, respectively (Table S7). Although the downstream orthologs lack confirmation, the regulation patterns in conifers seem different from angiosperms. However, the second highest scoring HSP for these genes showed similar patterns to the proposed model - simplified in [35]. That is, UNDEAD-CJt088123 activated, whereas VGD1-CJt014021 and GLOX1-CJt035052 suppressed the gene expression level in LF (leaf tissue). It should be noted that Phan *et al*. (2011) [35] used young floral bud tissues as the control.

### Accessions specific variant events

For further applications for the sugi breeding program or genetic study, we mined over twenty-five thousands SNPs and indels of each group, which were assigned to seven parental lines in Table S8. They all passed the customized filter - QUAL > 20 and DP > 3 (variant quality value above 20 and sequence depth of the variant site above 3). The abundance of the variants is based on counting the alternative alleles against the reference sequence. The variants has been uploaded in compressed VCF format on ForestGEN.

## Discussion

The CJ3006NRE sequence is the assembled product of the cDNA library integrated from 10 different libraries. We used two different approaches to integrate from each library’s assembled result. Then, we performed a second integration to unite the two integrated results into one. After eliminating potential contamination by BLAST against the NCBI-NR database, and filtering out potential assembly error by merging among libraries, the resulting sequence CJ3006NRE is suitable for investigating differential gene expression and structural annotations.

### Sequence depth is the one of the keys for accurate assembly

There is still some debate as to how to assemble the transcriptomic sequence data with high-quality. A greater sequencing depth or number of reads is a part of key in sampling all transcripts. In relation to the benchmark for S1s or S2s to S5s (Table1 and Fig 1), it is clear that a higher abundance of sequencing can cover the missing core genes, even if the abundance is not as high as S3s or S4s. Nevertheless, above some abundance, sequencing depth is no longer the limitation to increase the accuracy of assembly. For example, if we attempt to decrease missing parts to 15%, based on the number of reads and the BUSCO result, the optimal point would be at about 33 million read pairs (the equation for trend-line of the Missing in Figure S3). This number is not a perfect estimation; however it is a reasonable estimate from the trend.

### Comprehensive union

The condition of this study was that there were several libraries from three types of tissues: male-strobilus, leaf, and inner bark. However, these were collected from close but different genetic backgrounds, an asymmetrical number of runs, and abundances. Considering that alternative splicing happens among different conditions, e.g., tissues or treatment, we independently assembled each library, and then joined them to an integrated library. In assembly with Trinity, the assembled number of contigs ranged from 43,326 (S5) to 124,448 (S3s) for each library (Table 1). The number of assembled contigs was about the same as that by Oases (Table S1). We used EvidentialGene to integrate these 50 assemblies, including 10 by the Trinity and 40 by the Oases, into one library of 108,886 contigs. The second set consisted of 40,368 contigs from 10 libraries, where the data were manually integrated and assembled in Trinity (Figure S4). Using EvidentialGene as well as manually integration, the 49,758 sugi CJ3006NRE has resulted in high confidence in the sequencing dataset.

Repetitive sequences and isoforms co-exist in CJ3006NRE. Their compositions reflect the variation in the conditions of the samples collected. As for now (as it was in the pre-genomic era of sugi), increases in accuracy and efforts toward completeness of the transcriptome data is important. On the other hand, the redundancy has been dealt with using two separate methods, EvidentialGene and manual integration. There are two advantages from integrate-after-assembly: one is that the isoform can be kept, the other one is that they could be used to validate contigs among libraries. It is obvious that if the assembly tools have a lower miss-assembly rate, then we could keep the isoform for subsequent structural annotation, while the pseudomolecules are available. The latter is to overcome the miss-assembly due to randomized k-mer alignment processes. Because we only kept contigs which existed multiply among libraries, chance of producing the same miss-assembled contig would be low, but the isoforms or gene family may be a part of the miss-assembly.

### Discarding contamination

Filtering out contamination is an important step in reducing redundancy. CJ3006NRE was selected based on the alignment results against NCBI-NR and the taxa belonging to “Eukaryote”. Although algae and fungi could also be collected while sampling from sugi tissue, the term “Eukaryote” may be too loose to filter out contamination. “Euphyllophyta” and “Spermatophyta” are the other potential thresholds, as they are more specific. However, considering that the accumulation of genomic data for conifers has not been as established as it has been in the angiosperms, orthologs in sugi may have less homology that could be identified by BLAST. We used the “not Eukaryote” term as our negative threshold in order to discard the “non-Eukaryota” types, including those in Archaea, Bacteria, etc. Surprisingly, the reduction in the mapping rate of input reads was only less than 4%. Thus, we suggest the use of CJ3006NRE as the reference transcriptome, in place of CJ3006All.

### Increasing the accuracy of assembly

Based on the benchmark results, completeness, especially “single-copy,” increased to up to 78.47% of CJ3006NRE from less than 45% of the S4s library. This suggests the integration process dramatically decreased the duplication and fragments. Following the ideas of BUSCO [34], a higher number of duplicates may indicate a more erroneous assembly of haplotype. Thus, according to the evaluation result using BUSCO, our assembled contig has less duplicates. So our manipulation effectively increased the quality of the assembly.

The more full-length cDNA is available, the higher the accuracy for subsequent annotations on the genomic sequences. We used both third-generation sequencing and full-length cDNA library downloaded from NCBI as the validation resource. In the alternative method of classical PCR procedures, primers would have to be designed and the labor costs of bench work would be enormous. If we take ISOSeq0215 (56,399 sequences) as the validation standard and above 90% coverage of genes as the threshold, CJ3006NRE could cover 35,741 (63.3%) ISOSeq0215 sequences by using only 12,231 (24.6 %) CJ3006NRE out of 49,758 (Table S9). Thus, we could learn two points from here. The first, about 25% of CJ3006NRE could be validated by ISOSeq0215 at least. The second, by the statistics, we found there were more CJ3006NRE with the potential full-length cDNA than the other cDNA resources.

Overall, these results indicate that the methodology used in this study would be useful for assembling transcriptome data of non-model plant organisms.

### Contig number and all sugi genes

The total number of loci in sugi haplotype is uncertain without genomic sequencing. Since we have the total RNA from male strobili, leaf, and bark, studies on other tissue types and conditions will be fruitful with annotation as the goal. However, if we compare to two model angiosperm species, *Oryza sativa* [36, 37] and *Arabidopsis thaliana* [38, 39], our result may be better than the BUSCO benchmark estimation. Considering that these two model plant has been extensively annotated and continuously updated for over a decade, both have fine genomic sequences. Without discarding the transposable elements (TE), the loci numbers for rice and *Arabidopsis* are 37,848 and 38,194, respectively. In CJ3006NRE, if we discard non-representative isoforms, the gene number of with or without TE are 39,762 and 36,947, respectively. The total gene number is similar to these two model plants.

Of course, the discovery of more functional genes from collecting different transcriptome data from different tissues or conditions is expected in future. As BUSCO estimated (Figure 1), about 10.7% of the core genome has not been identified yet. Part of the 10.7% may include differences between the core genome of angiosperms and gymnosperms. This may become clearer with the development of more advanced sequencing technology and more gymnosperms genomic data becoming available.

The gene number in sugi was higher than the two model plants. Within the 49,758 contigs, 487 (0.98%) genes were not TE and cannot be annotated by InterProScan or EvidentialGene. Similar sequences were also not found in the NCBI-NR database, with an e-value = 10. The length of these unidentified genes ranged from 382 bp to 5507 bp. The protein length by these genes ranges from 18 to 197 amino acids. Thus, they have UTR and CDS regions that could be translated into protein sequences. The 487 non-TE genes may be potentially novel genes that could provide the new insights of functional genomics in conifers.

### Different composition of transcription factors

Because the gene regulatory network represents the ordering of gene expression, understanding the transcription factors is the key to understand the pathways and the mechanisms involved in metabolism. In addition, as these transcription factors play a key role in the fundamental pathways (e.g., the MYB gene family)[40, 41], the highly conserved domain and functionally conserved fragments reduce the potential for misidentification of orthologues. Since we attained about 90% of the sugi total genome assessed by BUSCO, the transcription factors we found should have approximately the same coverage to all transcription factors in the sugi genome. The classification result of sugi and the two model plants is listed in Table S10. Compared to *Arabidopsis* and rice, the total number of transcription factors in sugi is 1,340, which is less than two angiosperms (1,924 and 1,455, respectively). The CSD (cold-shock domain) is the one of the lowest in sugi, whose functions, under the name “cold-shock”, has been identified to help the cell to survive under low temperature conditions. Surprisingly, only one has been found in our samples. Sugi might have some different strategies to tolerate cold conditions than the other two model species since the life cycle is long and perennial. This may represent different strategies used to tolerate cold temperatures among conifers and angiosperms; otherwise they may have not been expressed during the collection of mRNA. In the other categories, some trends and numbers of transcription factors between sugi and the two angiosperms were similar to that of a previous study [42], but not always. For example, although the number of transcription factors is not as low as in maritime pine for “zf-Dof”(PF02701) transcription factors [42], it was still the smallest in sugi. This number is the same as in the model moss *Physcomitrella patens* [43]. Only 20 genes with zf-Dof in sugi were identified in our study.

The number of transcription factors may reflect the complexity of the regulatory networks. Especially there are differences in the scale of environmental changes over the whole life cycle as well as the lifespan. Sugi trees could have a lifespan of over many years, but *Arabidopsis* lives for around two months. However, the influence of the functional domain of a protein sequence is the most important of all the features even between distant species. In previous studies of several MYB gene families or categories, e.g., MYB80 [35, 44] and 3R-MYB [45], the conserved sequences are sometimes the key to normal function (e.g., metabolic), and could be used as a footage to assess evolution in lineages.

### Tissue-specific gene expression

Within the 7,776 significant differences in gene expression between the male strobili and leaf materials, the MF (expression level higher in male strobili) group contains more genes than the IBL (expression level higher in inner bark or leaf) group (4,471 vs. 3,305). Comprehensive viewing of the distribution via gene ontology (GO) showed no obvious differences among the categories (Table S11). However, the pattern of gene expression showed significant differences based on the PCA analysis (Fig 3a).

We focused on transcription factors to further explore certain categories of genes. In the MF group, the three MYB genes with highest expression level and significant difference against LF were CJt069044, CJt099797, and CJt070090. Based on the phylogenetic analysis (Figure S5), the first two are the orthologs of AtMYB35 (TDF1; AT3G28470) and AtMYB80 (AT5G56110). In turn, they are considered to be highly expressed in the anthers and play the important role in tapetum development [35, 46]; and thus can also determine whether male sterility occurs. Our results revealed that these two pairs of orthologs were highly expressed in the similar organs, in the anthers of angiosperms and the male strobili of conifers. Although there was no validation from orthologs, the potential orthologs of genes downstream from AtMYB80 nonetheless showed similar trends in the male strobili (Figure 3c). This suggests the genes downstream of MYB80 are homologous in angiosperms and gymnosperms. CJt070090 in sugi is a 3R-MYB. The most similar gene in *Arabidopsis* is AtMYB3R4 (AT5G11510), which was highly expressed in anther and young leaf tissues (based on the “BAR eFP Browser” in TAIR website). It has been suggested that the gene may be involved in suppressing mitosis [47]. However, CJt070090 is mostly not expressed in leaf tissues (Figure 3c). Although we cannot be sure about the evolutionary relationships between CJt070090 and AtMYB3R4, either they are orthologous or out-paralogous; the expression trend in different organs is the same as for CJt099797 (Figure 3c). It means both CJt099797 and CJt070090 are only expressed in male strobili, but not in leaves. This suggests CJt070090 may play more specific role in the male strobili of sugi in comparison to AtMYB3R4 in anther and young leaf tissues of *Arabidopsis*.

## Conclusion

In this work, we performed *de novo* assembly of *Cryptomeria japonica*, sugi, by integrating transcriptome data from unequal runs of 10 libraries. They were collected from two different types of tissues with a slightly different genetic background within a short period. By using the public pipeline, EvidentialGene and half-manual integration, we balanced controlling miss-assembly and wasting too many raw reads. According to the pedigree of the samples, the potential SNPs and indels recognized in this study could be useful for future breeding or genetic research. The high confidence of the contigs and translated protein sequences is a novel resource for the conifer study community. Furthermore, evolutionary studies will be benefited by the additional gymnosperm genomic data.

## Materials and methods

### Plant materials

Ten accessions were prepared for mapping the male-sterile gene. The pedigree is shown in Figure S1. ‘Nakakubuki-4’ is the male parent of ‘T1NK4F1’. T5_normalMIX_ms1 and T5_sterileMIX_ms1 are from the progeny of ‘T1NK4F1’ backcrossed with ‘Toyama MS’, and are the male fertile *Ms1*/*ms1* and male sterile *ms1*/*ms1*, respectively. These four accessions will be called T5 family hereafter. S3T67_normalMIX_ms1 and S3T67_sterileMIX_ms1 are samples from the progeny crossed with ‘T1NK4F1’ (male fertile) and ‘Shindai-3’ (male sterile), respectively. ‘Ooi-7’ carries a heterozygous male-sterile gene (*Ms1*/*ms1*). ‘S1NK4’ is the F1 hybrid of ‘Shindai-1’ and ‘Nakakubuki-4’. ‘S5HK7’ is the F1 hybrid of ‘Shindai-5’ and ‘Higashikanbara-7’. ‘S8HK5’ is the F1 hybrid of ‘Shindai-8’ and ‘Higashikanbara-5’. Most parental lines of these crosses above are not included in this study, except for ‘Nakakubuki-4’; but they provided four different male-sterile genes. Since the genetic characters of these male-sterile genes have been well studied, they are all recognized to be recessive genes. In these samples, only T5_sterileMIX_ms1 and S3T67_sterileMIX_ms1 are clearly male-sterile groups.

### Extraction of RNA and sequencing

The 10 RNA-Seq libraries were constructed from 10 different accessions. For S3s, S4s, S5s, and S6s, RNA was extracted from several individuals (up to 50 individuals) and the RNA mixture of progeny was sequenced. RNA-Seq for S1s to S6s was carried out, as described in [19]. For the rest of the libraries, we extracted RNA from different tissues (Table 1) following the method used in [19], and the mixture of RNA from these tissues was sequenced on HiSeq2000 (Hokkaido System Science Co., Ltd, Sapporo, Japan). We performed ISO-Seq using the RNA of S2s and sequenced on PacBio RS (Takara Bio Inc.) with four cells by P3C5 chemistry.

### Assembly and annotation

#### Quality control and independent assembly

Before assembly, the raw reads were passed through four quality controls using Cutadapt [48]. These included: (1) cutting 13 bases from five prime sides, (2) cutting the over-reading due to adapters or primers, (3) cutting the low quality base tails, and (4) setting the minimum length threshold to 35 bases after the other steps. The filtered numbers of reads are listed in Table 1. Hereafter, “library” means the union of one or multiple runs of RNA-Seq data from the same accession or variety.

All 10 libraries were assembled using two different software - Oases, v0.2.08 [29] and Trinity v 2.4.0 [30]. Since the maximum k-mer for Trinity was 32, we performed only a single run for each library. For Oases (Velvet), we used the k-mer in the range from 35 to 43 (odd numbers), with five runs for each library. For both software, the minimum length of contigs was 500 and 300 for Trinity and Oases, respectively. The minimum length for Oases has considered the existence of non-coding RNA.

#### Integration

Contig integration was used to overcome the bias from sampling to assembly. We used two integration methods to increase the reliability of integrated contigs (Figure S4). One method was automatic pipeline using EvidentialGene [28], which is open source software. The other is half-manual pipeline using homemade scripts to manipulate the assembled result from Trinity. The coding languages used include Shell script, AWK, and Python.

In the workflow through EvidentialGene [28], each library was processed by tr2aacds.pl once to produce 10 independently integrated contigs (FASTA files). This step was intended to keep the isoforms that were produced under different conditions. Then, the 10 contig FASTA files were concatenated into one and again processed by tr2aacds.pl. The only customized parameter was the setting for the minimum length of CDS with 90, ie. “--MINCDS90”.

After the second integration using EvidentialGene, there were 107,674 transcripts which could be converted into 108,886 protein sequences.

To begin the half-manual integration, the assembled contigs have been translated to the longest open reading frame as the represented protein sequences.

The half-manual integration process includes three modules: 1) finding the seed sequence, 2) mining more paralogs, and 3) abstracting by homologs; named module-1, module-2, and module-3, respectively. Both nucleotide and protein sequences could be used as an operating object, depending on whether the alignment within species or among species.

In module-1, the main objective was to pick seed sequences via BLASTX. We expected these seeds contained the functional domain. These do not necessarily have to be orthologs of the subject. We used SwissProt [49] (downloaded June 11, 2017) as a reference to find the orthologous genes for each library. Two filters were applied to the BLASTX result: 1) Minimum ratio of the sum of HSP against the subject’s sequence was 50%; 2) Minimum HSP length was 20 amino acids. The contigs with the best score were selected for every SwissProt gene. Thus, one SwissProt gene was only linked to one contig. On the contrary, after a backward filtering, one contig was only linked to one SwissProt gene. After this step, we obtained one set of cDNA sequence used as the “seed” sequence for the next step.

In module-2, we aimed to extend and concentrate by identifying paralogs. The similar genes have been identified in module-1 as seed genes, which represented the most similar genes across angiosperms and gymnosperms. Any duplication after the separation of angiosperms and gymnosperms, which produced the paralogs, could be identified from the seed genes in this step. The “extend” query used the seed sequences to fish for more paralogs. We used “seed” sequences as a query against all 10 libraries, including the source library of the “seed”. There were no customized conditions for running the BLAST on this step. In this step, since the contigs were from the same species, we expected to retrieve as many homologous contigs in the sequence as possible. The “concentrate” query was used to reduce the duplication sequences to be as representative as possible. To find the representative sequence among homologs, the maximum tolerance for mismatches and gaps was 2 bps. However, we only kept the representative sequence that could be found similar enough and existed at least two libraries.

The last step in module-2 was to include the correct (“good”) singletons. Since the previous “extent” and “concentrate” steps resulted in singletons, we passed the filter “Find seed sequences”, but did not find homologs in any other library. These “good” singletons have to be added back in the next step.

In the last step, module-3, by referring to the taxonomic information in the NCBI-NR database, we discarded those that matched to non-eukaryotes, but kept those that matched to eukaryotes and others to unclassified.

#### Annotation

We used two tools, EvidentialGene [28], and InterProScan (v 5.30) [31], to annotate the integrated library (CJ3006NRE). The reference databases for running namegenes.pl, the annotation tool in EvidentialGene, were UniRef50 (downloaded May 2018) [50] and CDD (Conserved Domains Database, v 3.16). The reference database used for annotation was InterPro5 v 69.0.

The isoforms would be identified using BLAST against all contigs with a parameter, “word size = 100 bps.” Then, the contig was matched to another contig with over 90% of genes identified. An HSP length of over 150 bps would be considered as an isoform.

For identifying the transcription factors of sugi, we re-scanned all the contigs using pfamScan [32]. The list of transcription factors was based on a joint list with the work of [42] and a Pfam list published online, www.transcriptionfactor.org [51].

For additional prediction of metabolic enzymes, we used a pipeline called E2P2, downloaded from the “Plant Metabolic Network” (PMN) [52].

In order to mark the repetitive sequences, RepeatMarker v 4.0.7 (http://www.repeatmasker.org) [33] and RepBase v 22.05 were used as the reference databases.

#### Evaluation

For evaluating the proportion of input reads that have been wasted during integration, we aligned the input reads, as done for assembling, to the originally assembled contigs and the integrated one. We used BWA [53, 54] as mapping tools. Considering that *Cryptomeria japonic*a is a heterogeneous species, we tuned down the parameter of penalty. For BWA, we used “bwa mem” module and one of the parameters we set was “-O 4,4”, representing penalties for deletions and insertions, which means gaps on reads and on references, respectively.

For estimating the coverage of the core orthologous genes, we used BUSCO (v 3.0.2) [34] for testing all 10 libraries and the integrated library. The testing model was “transcriptome” and the version of the reference database was “embrophyta_odb9”. For each of 10 RNA-Seq libraries, we used their own assembled contig as the input; for the integrated library, we used the CJ3006NRE sequence as the input.

The ISO-Seq data were processed using the “pbtranscript-tofu” analysis suite v 1.0.0.177900 [55]. The only customized parameter was 300 bps as the minimum length; the rest were default values.

A total of 23,111 full-length cDNA sequences were downloaded from NCBI. We retrieved these sequences by using the keywords, “Cryptomeria japonica”, “full-length”, and “cDNA” using the NCBI web-based searching interface on 13^th^ September, 2018..

### Differential expression

Using our integrated library, CJ3006NRE, as a reference, we compared the expression levels among the 10 libraries. “Kallisto” [56] was used to quantify the transcript abundances using bootstrap estimation from 100 repetitions. “Kallisto” is a package which calculates the building index of the reference sequence and quantifies the abundance from FASTQ files. The output of “Kallisto” was processed in differential expression analysis using “sleuth.”

### Gene Ontology (GO) annotation

The GO terms were assigned with InterProScan during the annotation process. The classification of the GO terms was done using CateGOrizer [57] using Plant_GOslim as the classification list.

### Variant calling

Upon obtaining the mapped files—BAM (binary SAM), we used samtools [58] and bcftools (https://github.com/samtools/BCFtools) to call the variants. The group-specific variants were classified using the “isec” command in bcftools. Appendix S1 shows an example of command lines. Table S8 summarized the variantions among the accessions against CJ3006NRE cDNA set. The variant-calling done by bcftools. According to this Table and pedigree (Figure S1), we extracted the group (or accession) specific variant. E.g., for ‘Shindai 3’, there was only one site with allele “1” in S5 and S6s, and allele “1” was not present in the rest of the sites (Figure S6).

## Accession numbers

Sequences used in the present study have accession numbers DRR174638 to DRR174656.

## Acknowledgements

The authors would like to thank Shinji Itoo for producing a part of the materials used in the current study. Arboretum and Nursery Office, FFPRI was involved in the maintenance of *C. japonica* nursery. Computations were mostly performed on the supercomputer of AFFRIT, MAFF, Japan. This work was partly supported by a Grant-in-Aid from the Program for Promotion of Basic and Applied Researches for Innovations in Bio-oriented Industry (No.28013B), JSPS KAKENHI (Grant Number 25450223), Research grants #201119 and #201421 from the Forestry and Forest Products Research Institute, and grants from the Project of the NARO Bio-oriented Technology Research Advancement Institution (Research program on development of innovative technology (No.28013BC)).

## Supporting Information Captions

Table S1. Statistics of assembled libraries.

Table S2. The annotation of CJ3006NRE

Table S3. Major components of the transposable elements within CJ3006NRE and other libraries (a) and RepeatMasker output for CJ3006NRE (b)

Table S4. The mapping rate of raw reads against different reference sequences

Table S5. Differentially expressed transcription factors between “male strobilus” (MF) and “inner bark and leaf” (IBL) tissue set

Table S3. Gene families and number of transcription factors in sugi showing significant differences between male strobius (MF) and inner bark and leaf (IBL).

Table S7. Differential expression of MYB80 related genes of male strobili (MF) against inner bark and leaf (IBK)

Table S4. Specific variations of each parental line

Table S5. Mutual number of sequences (CJ3006NRE/cDNA source) under different coverage threshold

Table S10. Number of transcription factors in CJ3006NRE and other two model plants

Table S11. Number of gene ontology annotations for differentially expressed CJ3006NRE genes between male strobili (MF) and inner bark and leaf (IBL) sample

Figure S1. The pedigree of accessions.

Figure S2. Heatmap of significant differences in expressed genes among different tissue types.

Figure S3. The read number against the BUSCO benchmark result.

Figure S4. Workflow of assembly: a.) The general workflow of assembly; b.) The workflow of half-manual assembly

Figure S5. Phylogenetic tree for classifying three sugi MYBs.

Figure S6. Diagram for identifying group-specific variants.

Appendix S1 Example of command lines for variant calling.

## Notes

### Competing Interest Statement

The authors have declared no competing interest.

## References

1. Forestryagency M (2018) Annual Report on Forest and Forestry in Japan. Forestry Agency, Ministry of Agriculture, Forestry and Fisheries, Japan.

2. Saito Y (2014) Japanese cedar pollinosis: Discovery, nomenclature, and epidemiological trends. Proceedings of the Japan Academy, Series B 90: 203–210.

3. Otake T (2017) When pollen attacks! Experts reveal new approaches to combating hay fever. Japan Times.

4. Saito M (2010) Breeding Strategy for the Pollinosis Preventive Cultivars of Cryptomeria japonica D. Don. Journal of the Japanese Forest Society 92: 316–323.

5. Moriguchi Y, Uchiyama K, Ueno S, Ujino-Ihara T, Matsumoto A, et al. (2016) A high-density linkage map with 2560 markers and its application for the localization of the male-sterile genes *ms3* and *ms4* in *Cryptomeria japonica* D. Don. Tree Genetics & Genomes 12.

6. Mishima K, Hirao T, Tsubomura M, Tamura M, Kurita M, et al. (2018) Identification of novel putative causative genes and genetic marker for male sterility in Japanese cedar (*Cryptomeria japonica* D.Don). BMC Genomics 19.

7. Tsubomura M, Kurita M, Watanabe A (2016) Determination of male strobilus developmental stages by cytological and gene expression analyses in Japanese cedar (*Cryptomeria japonica*). Tree Physiology 36: 653–666.

8. Chen J, Uebbing S, Gyllenstrand N, Lagercrantz U, Lascoux M, et al. (2012) Sequencing of the needle transcriptome from Norway spruce (*Picea abies* Karst L.) reveals lower substitution rates, but similar selective constraints in gymnosperms and angiosperms. BMC Genomics 13: 589.

9. Birol I, Raymond A, Jackman SD, Pleasance S, Coope R, et al. (2013) Assembling the 20 Gb white spruce (*Picea glauca*) genome from whole-genome shotgun sequencing data. Bioinformatics 29: 1492–1497.

10. Warren RL, Keeling CI, Yuen MMS, Raymond A, Taylor GA, et al. (2015) Improved white spruce (*Picea glauca*) genome assemblies and annotation of large gene families of conifer terpenoid and phenolic defense metabolism. The Plant Journal 83: 189–212.

11. Zimin A, Stevens KA, Crepeau MW, Holtz-Morris A, Koriabine M, et al. (2014) Sequencing and Assembly of the 22-Gb Loblolly Pine Genome. Genetics 196: 875.

12. Guan R, Zhao Y, Zhang H, Fan G, Liu X, et al. (2016) Draft genome of the living fossil *Ginkgo biloba*. GigaScience 5.

13. Hizume M, Kondo T, Shibata F, Ishizuka R (2001) Flow Cytometric Determination of Genome Size in the Taxodiaceae, Cupressaceae *sensu stricto* and Sciadopityaceae. CYTOLOGIA 66: 307–311.

14. Nystedt B, Street NR, Wetterbom A, Zuccolo A, Lin YC, et al. (2013) The Norway spruce genome sequence and conifer genome evolution. Nature 497: 579–584.

15. Campbell MS, Holt C, Moore B, Yandell M (2014) Genome Annotation and Curation Using MAKER and MAKER-P. Current Protocols in Bioinformatics 48: 4.11.11-14.11.39.

16. Canas RA, Li Z, Pascual MB, Castro-Rodriguez V, Avila C, et al. (2017) The gene expression landscape of pine seedling tissues. Plant J 91: 1064–1087.

17. Carrasco A, Wegrzyn JL, Durán R, Fernández M, Donoso A, et al. (2017) Expression profiling in Pinus radiata infected with Fusarium circinatum. Tree Genetics & Genomes 13.

18. Rigault P, Boyle B, Lepage P, Cooke JEK, Bousquet J, et al. (2011) A White Spruce Gene Catalog for Conifer Genome Analyses. Plant Physiology 157: 14–28.

19. Ueno S, Kentaro U, Moriguchi Y, Ujino-Ihara T, Matsumoto A, et al. (2019) Scanning RNA-Seq and RAD-Seq approach to develop SNP markers closely linked to *MALE STERILITY 1 (MS1*) in *Cryptomeria japonica* D. Don. Breeding Science.

20. Xiang X, Zhang Z, Wang Z, Zhang X, Wu G (2015) Transcriptome sequencing and development of EST-SSR markers in Pinus dabeshanensis, an endangered conifer endemic to China. Molecular Breeding 35.

21. Brenner ED, Katari MS, Stevenson DW, Rudd SA, Douglas AW, et al. (2005) EST analysis in *Ginkgo biloba*: an assessment of conserved developmental regulators and gymnosperm specific genes. BMC Genomics 6: 143.

22. Futamura N, Totoki Y, Toyoda A, Igasaki T, Nanjo T, et al. (2008) Characterization of expressed sequence tags from a full-length enriched cDNA library of Cryptomeria japonica male strobili. BMC Genomics 9: 383.

23. Du S, Sang Y, Liu X, Xing S, Li J, et al. (2016) Transcriptome profile analysis from different sex types of *Ginkgo biloba* L. Frontiers in Plant Science 7.

24. Abdel-Ghany SE, Hamilton M, Jacobi JL, Ngam P, Devitt N, et al. (2016) A survey of the sorghum transcriptome using single-molecule long reads. Nature Communications 7: 11706.

25. Zimin AV, Puiu D, Luo MC, Zhu T, Koren S, et al. (2017) Hybrid assembly of the large and highly repetitive genome of Aegilops tauschii, a progenitor of bread wheat, with the MaSuRCA mega-reads algorithm. Genome Res 27: 787–792.

26. Geniza M, Jaiswal P (2017) Tools for building de novo transcriptome assembly. Genomic resources and databases 11-12: 41-45.

27. Ueno S, Nakamura Y, Kobayashi M, Terashima S, Ishizuka W, et al. (2018) TodoFirGene: Developing Transcriptome Resources for Genetic Analysis of Abies sachalinensis. Plant Cell Physiol 59: 1276–1284.

28. Gilbert DG (2019) Longest protein, longest transcript or most expression, for accurate gene reconstruction of transcriptomes? bioRxiv: 829184.

29. Schulz MH, Zerbino DR, Vingron M, Birney E (2012) Oases: robust de novo RNA-seq assembly across the dynamic range of expression levels. Bioinformatics 28: 1086–1092.

30. Grabherr MG, Haas BJ, Yassour M, Levin JZ, Thompson DA, et al. (2011) Full-length transcriptome assembly from RNA-Seq data without a reference genome. Nat Biotechnol 29: 644–652.

31. Jones P, Binns D, Chang HY, Fraser M, Li W, et al. (2014) InterProScan 5: genome-scale protein function classification. Bioinformatics 30: 1236–1240.

32. Finn RD, Tate J, Mistry J, Coggill PC, Sammut SJ, et al. (2008) The Pfam protein families database. Nucleic Acids Res 36: D281–288.

33. Smit AFA, Hubley R, Green P (2013-2015) RepeatMasker Open-4.0.

34. Simao FA, Waterhouse RM, Ioannidis P, Kriventseva EV, Zdobnov EM (2015) BUSCO: assessing genome assembly and annotation completeness with single-copy orthologs. Bioinformatics 31: 3210–3212.

35. Phan HA, Iacuone S, Li SF, Parish RW (2011) The MYB80 transcription factor is required for pollen development and the regulation of tapetal programmed cell death in Arabidopsis thaliana. Plant Cell 23.

36. Tanaka T, Antonio BA, Kikuchi S, Matsumoto T, Nagamura Y, et al. (2008) The Rice Annotation Project Database (RAP-DB): 2008 update. Nucleic Acids Res 36: D1028–1033.

37. Kawahara Y, de la Bastide M, Hamilton JP, Kanamori H, McCombie WR, et al. (2013) Improvement of the *Oryza sativa* Nipponbare reference genome using next generation sequence and optical map data. Rice (N Y) 6: 4.

38. Huala E, Dickerman AW, Garcia-Hernandez M, Weems D, Reiser L, et al. (2001) The Arabidopsis Information Resource (TAIR): a comprehensive database and web-based information retrieval, analysis, and visualization system for a model plant. Nucleic Acids Research 29: 102–105.

39. Berardini TZ, Reiser L, Li D, Mezheritsky Y, Muller R, et al. (2015) The arabidopsis information resource: Making and mining the “gold standard” annotated reference plant genome. genesis 53: 474–485.

40. Altay G, Emmert-Streib F (2010) Inferring the conservative causal core of gene regulatory networks. BMC Systems Biology 4: 132.

41. Strygina KV, Khlestkina EK (2019) Myc-like transcriptional factors in wheat: structural and functional organization of the subfamily I members. BMC Plant Biology 19: 50.

42. Canales J, Bautista R, Label P, Gómez-Maldonado J, Lesur I, et al. (2014) De novo assembly of maritime pine transcriptome: implications for forest breeding and biotechnology. Plant Biotechnology Journal 12: 286–299.

43. Jin J, Tian F, Yang D-C, Meng Y-Q, Kong L, et al. (2016) PlantTFDB 4.0: toward a central hub for transcription factors and regulatory interactions in plants. Nucleic Acids Research 45: D1040–D1045.

44. Browne RG, Iacuone S, Li SF, Dolferus R, Parish RW (2018) Anther Morphological Development and Stage Determination in Triticum aestivum. Frontiers in Plant Science 9: 228.

45. Feng G, Burleigh JG, Braun EL, Mei W, Barbazuk WB (2017) Evolution of the 3R-MYB Gene Family in Plants. Genome Biology and Evolution 9: 1013–1029.

46. Zhu J, Chen H, Li H, Gao J-F, Jiang H, et al. (2008) Defective in Tapetal Development and Function 1 is essential for anther development and tapetal function for microspore maturation in Arabidopsis. The Plant Journal 55: 266–277.

47. Chandran D, Inada N, Hather G, Kleindt CK, Wildermuth MC (2010) Laser microdissection of <em>Arabidopsis</em> cells at the powdery mildew infection site reveals site-specific processes and regulators. Proceedings of the National Academy of Sciences 107: 460.

48. Martin M (2011) Cutadapt removes adapter sequences from high-throughput sequencing reads. EMBnetjournal; Vol 17, No 1: Next Generation Sequencing Data Analysis.

49. Bairoch A, Apweiler R (1996) The SWISS-PROT protein sequence data bank and its new supplement TREMBL. Nucleic Acids Res 24: 21–25.

50. Suzek BE, Wang Y, Huang H, McGarvey PB, Wu CH, et al. (2015) UniRef clusters: a comprehensive and scalable alternative for improving sequence similarity searches. Bioinformatics 31: 926–932.

51. Kummerfeld SK, Teichmann SA (2006) DBD: a transcription factor prediction database. Nucleic Acids Research 34: D74–D81.

52. Schläpfer P, Zhang P, Wang C, Kim T, Banf M, et al. (2017) Genome-Wide Prediction of Metabolic Enzymes, Pathways, and Gene Clusters in Plants. Plant Physiology 173: 2041.

53. Li H, Durbin R (2010) Fast and accurate long-read alignment with Burrows-Wheeler transform. Bioinformatics 26: 589–595.

54. Li H, Durbin R (2009) Fast and accurate short read alignment with Burrows-Wheeler transform. Bioinformatics 25: 1754–1760.

55. Gordon SP, Tseng E, Salamov A, Zhang J, Meng X, et al. (2015) Widespread Polycistronic Transcripts in Fungi Revealed by Single-Molecule mRNA Sequencing. PLOS ONE 10: e0132628.

56. Pimentel HJ, Bray N, Puente S, Melsted P, Pachter L (2016) Differential analysis of RNA-Seq incorporating quantification uncertainty. -.

57. Hu Z-L, Bao J, Reecy JM (2008) CateGOrizer: a web-based program to batch analyze gene ontology classification categories. Online Journal of Bioinformatics 9: 108–112.

58. Li H, Handsaker B, Wysoker A, Fennell T, Ruan J, et al. (2009) The Sequence Alignment/Map format and SAMtools. Bioinformatics 25: 2078–2079.

