## Supplemental figures and appendix for "Inspecting abundantly expressed genes in male strobili in sugi (*Cryptomeria japonica* D. Don) via a highly accurate cDNA assembly"

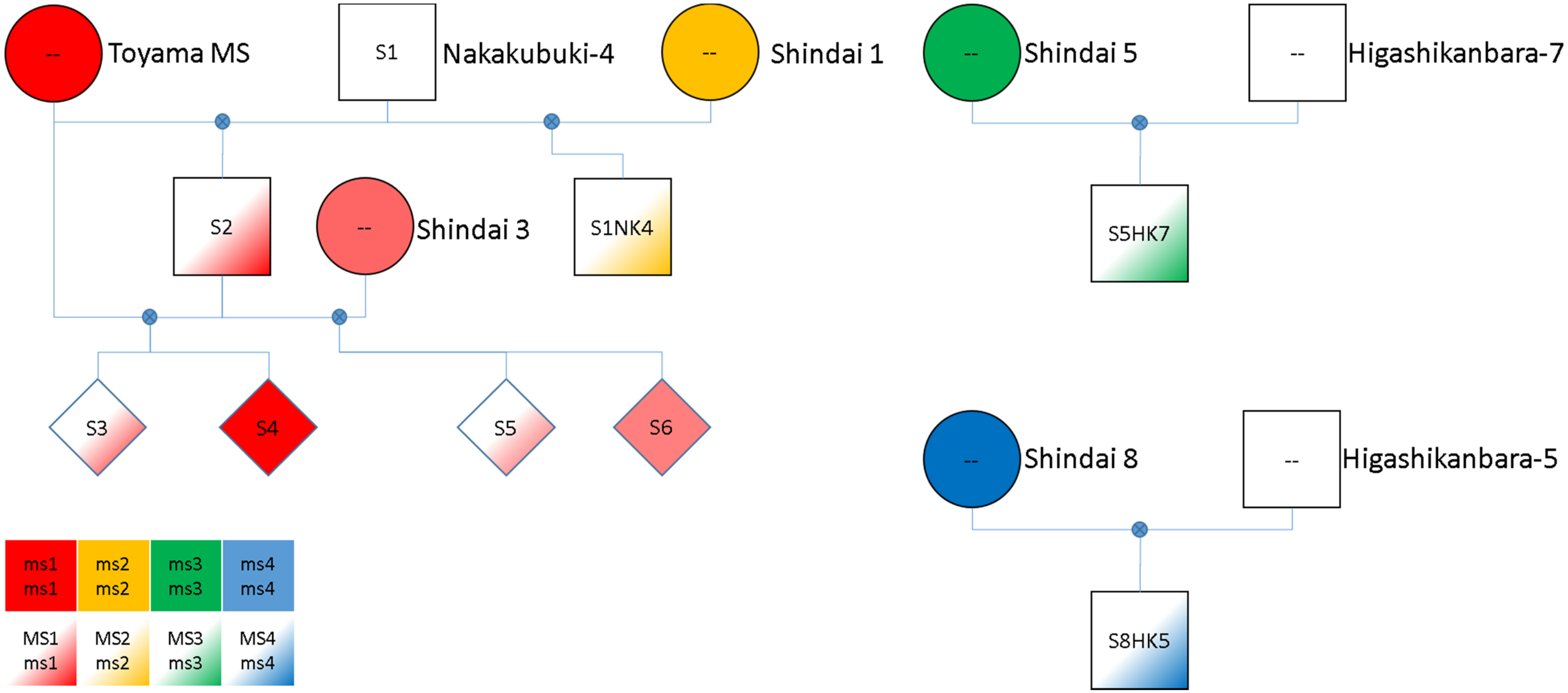


Figure S1. The pedigree of accessions.

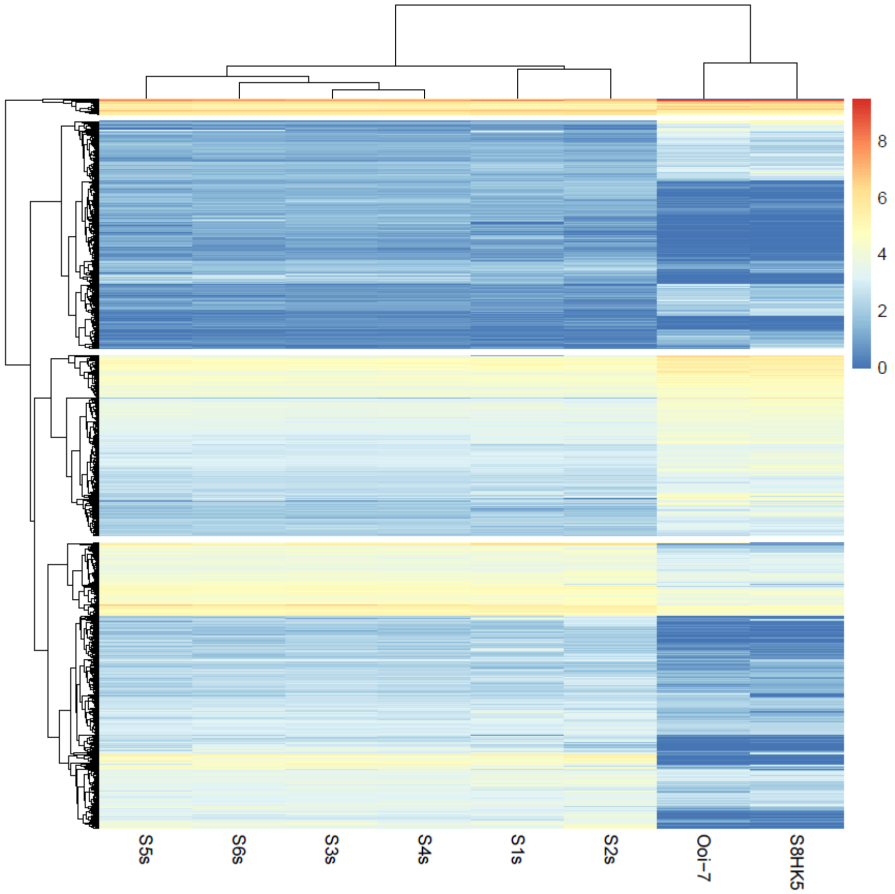


Figure S2. Heatmap of significant differences in expressed genes among different tissue types.

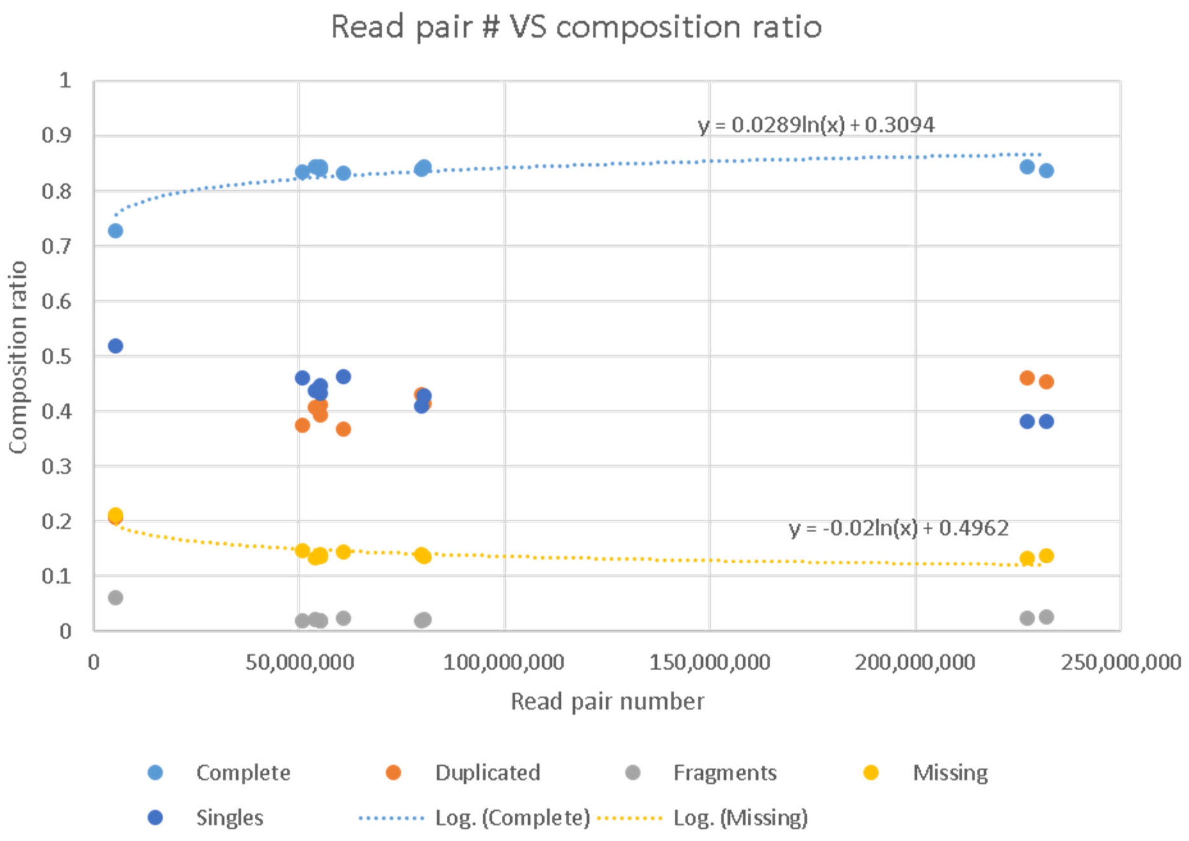


Figure S3. The read number against the BUSCO benchmark result.

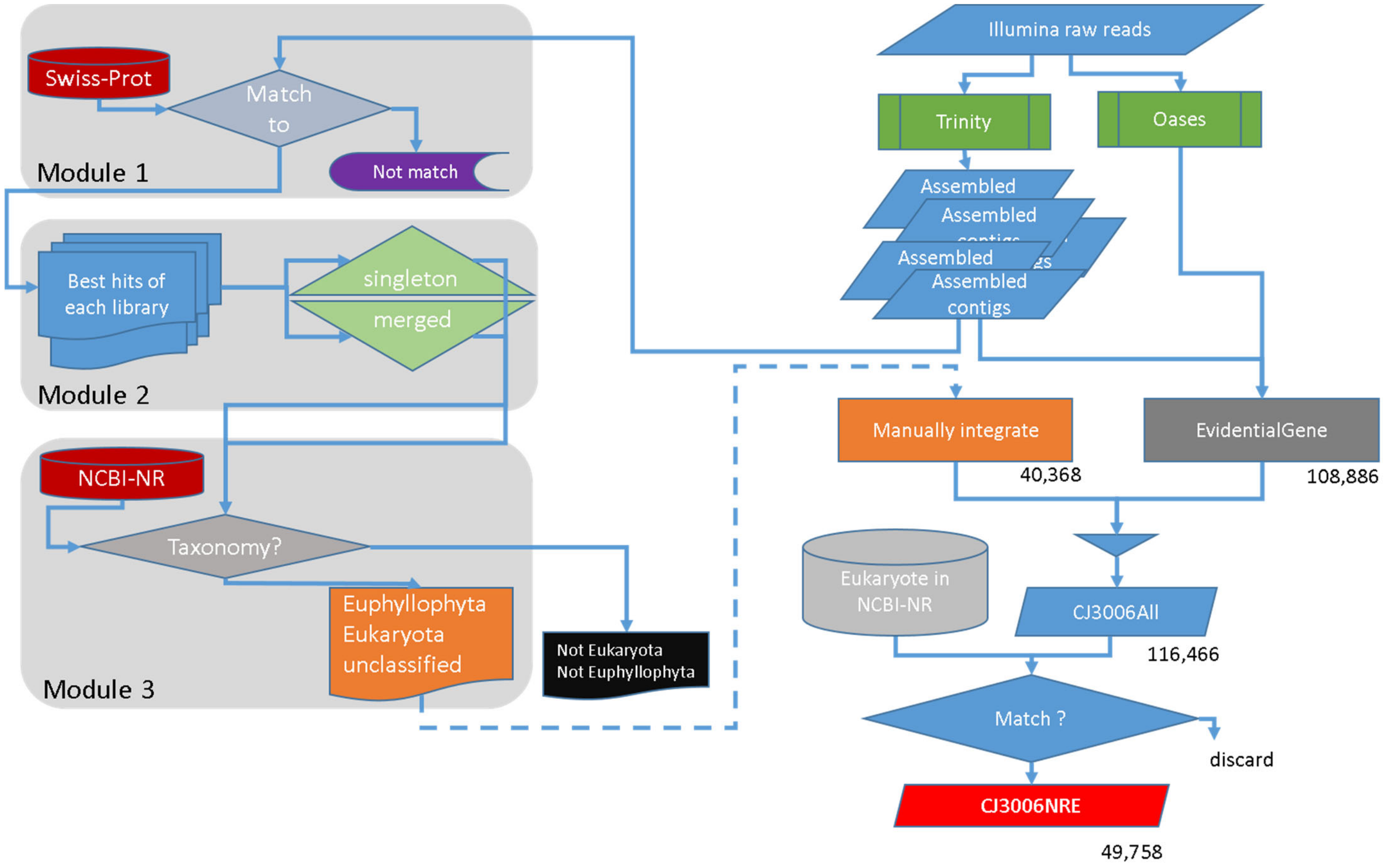


Figure S4. Workflow of assembly: a.) The general workflow of assembly; b.) The workflow of half-manual assembly

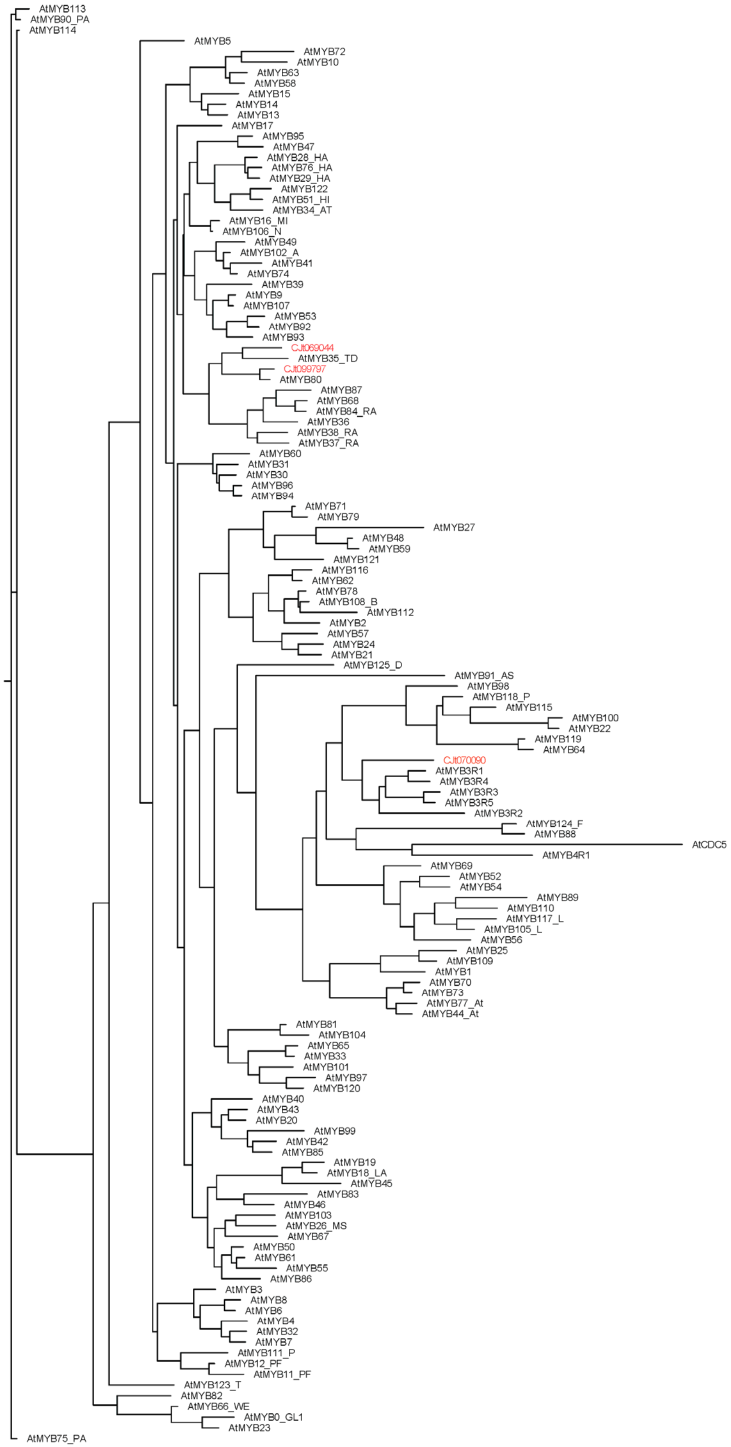


Figure S5. Phylogenetic tree for classifying three sugi MYBs.

| Specific for | S1s | S2s | S3s | S4s | S5 | S6s | Ooi-7 | S1NK4 | S5HK7 | S8HK5 |
|---|---|---|---|---|---|---|---|---|---|---|
| Nakakubiki-4 | 1 | 0 | 0 | 0 | 0 | 0 | 0 | 0 | 0 | 0 |
| Toyama MS | 0 | 1 | 1 | 1 | 1 | 1 | 0 | 0 | 0 | 0 |
| Shindai 3 | 0 | 0 | 0 | 0 | 1 | 1 | 0 | 0 | 0 | 0 |
| Shindai 1 | 0 | 0 | 0 | 0 | 0 | 0 | 0 | 1 | 0 | 0 |

Figure S6. Diagram for identifying group-specific variants.

```
#From BAM to VCFs
bcftools mpileup -L 1000 -m 3 -Ou -f CJ3006NRE.fa.gz S1s-CJ3006NRE.bam ¥
   | bcftools call -Amv > var_S1s-CJ3006NRE.raw.vcf
bcftools filter -s LowQual -e '%QUAL<20 || DP<3' var_S1s-CJ3006NRE.raw.vcf ¥
   | bgzip -c > var_S1s-CJ3006NRE.Q20DP3.vcf.gz
tabix -p vcf var_S1s-CJ3006NRE.Q20DP3.vcf.gz

#Find the intersection of any possible combination
bcftools isec var_S1s-CJ3006NRE.Q20DP3.vcf.gz ¥
    var_S2s-CJ3006NRE.Q20DP3.vcf.gz ¥
    var_S3s-CJ3006NRE.Q20DP3.vcf.gz ¥
    var_S4s-CJ3006NRE.Q20DP3.vcf.gz ¥
    var_S5s-CJ3006NRE.Q20DP3.vcf.gz ¥
    var_S6s-CJ3006NRE.Q20DP3.vcf.gz ¥
    var_Ooi-7-CJ3006NRE.Q20DP3.vcf.gz ¥
    var_S1NK4-CJ3006NRE.Q20DP3.vcf.gz ¥
    var_S5HK7-CJ3006NRE.Q20DP3.vcf.gz ¥
    var_S8HK5-CJ3006NRE.Q20DP3.vcf.gz -p ISEC/
```

Appendix S1 Example of command lines for variant calling.
